# ΦCrAss001, a member of the most abundant bacteriophage family in the human gut, infects *Bacteroides*

**DOI:** 10.1101/354837

**Authors:** Andrey N. Shkoporov, Ekaterina V. Khokhlova, C. Brian Fitzgerald, Stephen R. Stockdale, Lorraine A. Draper, R. Paul Ross, Colin Hill

**Affiliations:** APC Microbiome Ireland, University College Cork, Cork, Ireland

**Keywords:** crAssphage, crAss-like phages, human gut virome, bacteriophage, *Bacteroides*

## Abstract

ΦCrAss001, isolated from human faecal material, is the first member of the extensive crAssphage family to be grown in pure culture. The bacteriophage infects the human gut symbiont *Bacteroides intestinalis*, confirming *in silico* predictions of the likely host. Genome analysis demonstrated that the phage DNA is 102 kb in size, has an unusual genome organisation and does not possess any obvious genes for lysogeny. In addition, electron microscopy confirms that φcrAss001 has a podovirus-like morphology. Despite the absence of lysogeny genes, φcrAss001 replicates in a way that does not disrupt proliferation of the host bacterium and is able to maintain itself in continuous host culture.

---

The human gut virome contains a vast number of bacterial, mammalian, plant, fungal and archaeal viruses^1–3^. There is a growing body of evidence supporting specific and consistent alterations of the gut virome in a number of human diseases and conditions, including inflammatory bowel disease (IBD), malnutrition, and AIDS^4–6^. Currently, our understanding of the physiological significance of the human gut virome is limited by the fact that the vast majority of these viruses cannot be taxonomically classified or linked to any particular hosts (so-called “viral dark matter”^7^). Several recent studies identified a highly abundant (representing up to 90% of gut viruses in some individuals), unique uncultured group termed crAss-like phages or crAssphage^8–11^. Here we report on the first successful isolation of a crAssphage on a single host and describe its key biological properties. Based on electron microscopy of propagated crAssphage001 morphology we confirm that crAss-like phages possess podovirus-like morphology. We also demonstrate the ability of the phage to stably co-replicate with its *Bacteroides intestinalis* host in equilibrium for many generations *in vitro*, which mimics earlier observed ability of crAss-like phages to maintain stable colonization of the mammalian gut^11,12^.

In 2014, a study by Dutilh *et al*^8^ demonstrated the presence in ~50% of human samples of highly abundant sequences, which when cross-assembled from multiple sources, indicated the presence of a unique bacteriophage they named crAssphage (cross Assembly). The fully assembled 97 kb DNA genome showed no homology to any known virus, even though in some subjects it completely dominated the gut virome (up to 90% of sequencing reads). Indirect evidence suggested that *Bacteroides* species is a likely host^8^. A later study, based on detailed sequence analysis of proteins encoded by this phage, predicted *Podoviridae*-like morphology and placed it into the novel diverse and expansive family-level phylogenetic group of loosely related bacteriophages termed crAss-like phages^9^. Genomes of members of this group are highly abundant in the human gut and are also present in other diverse habitats including termite gut, terrestrial/groundwater, and oceans^9,13,14^. However, in the absence of a known host, no member of this family has been isolated and nothing is known of the biological properties of these crAss-like phages from the human gut.

In a recent analysis^11^, we *de novo* assembled 244 genomes of crAss-like phages from the human gut and classified them into several genus-and subfamily-level taxonomic groups based on percentage of shared orthologous genes. As a result of this study, we can state that 98-100% of healthy adults from Western cohorts carry at least one or more types of crAss-like phages, albeit with widely varying relative abundance. Here, we report the first documented isolation in culture of a crAss-like phage from the human gut virome and describe its key biological properties.

The replication strategy of crAss-like bacteriophages is unknown and all attempts to use standard plaque assays on semi-solid agar have failed. We therefore attempted to detect crAss-like phage replication using a broth enrichment strategy. Phage-enriched filtrates of faecal samples were collected from 20 healthy adult Irish volunteers, pooled and used to infect pure cultures of 53 bacterial strains representing the commensal human gut microbiota (Supplementary Table 1). After three successive rounds of enrichment, cell free supernatants were subjected to shotgun metagenomic sequencing. Analysis of the assembled sequencing reads demonstrated that the supernatant from strain *Bacteroides intestinalis* 919/174 was dominated by a single 102.7 kb contig (~98% of reads) related to a known but previously uncultured crAss-like bacteriophage, IAS virus, isolated from the human gut^9,15^. All other species of *Bacteroides* tested were insensitive to φcrAss001.

The genome of bacteriophage φcrAss001 is 102,679 bp and is circular or circularly permuted. We identified 105 protein coding genes (ORFs of length 90-7,239 bp) and 25 tRNA genes specific for 17 different amino acids. Cumulative G+C content of the phage genome is 34.7%, which is significantly lower than that of published *B. intestinalis* genomes (42.8-43.5%). An interesting structural feature of the genome is its apparent division into two parts of roughly equal size with strictly opposite gene orientation and inverted GC skew, possibly reflecting the direction of transcription and/or replication (Figure 1). Functional gene annotation was performed using a comprehensive approach, which included BLASTp amino acid sequence homology searches against NCBI nt database, hidden Markov model (HMM) searches against UniProtKB/TrEMBL database^16^ and profile-profile HMM searches with HHpred against PDB, PFAM, NCBI-CDD and TIGRFAM databases^17,18^. This allowed for the functional annotation of 57 genes and assignment of further 11 genes to conserved protein families with unknown functions. Ten identifiable structural protein genes (phage head, tail, appendages) as well as three genes responsible for lytic functions were clustered on the right-hand side of the genome, suggesting that the remaining un-annotated genes in this part of the genome may also be responsible for structure and assembly of phage particles, as well as cell lysis. By contrast, the left-hand side predominantly harboured genes involved in replication, recombination, transcription and nucleotide metabolism. Putative DNA-binding and transcriptional regulation proteins were located in the proximal portions of the two oppositely oriented genome halves, suggesting their role in governing the transcription of gene modules located downstream on both sides of the genome (Fig. 1). Two genes (3, β-fructosidase and 5, ferredoxin-thioredoxin reductase) were predicted to be involved in auxiliary metabolic processes, in that they are unrelated with phage replication and virion assembly. No identifiable lysogeny module, or integrase or recombinase genes were identified. MALDI-TOF analysis of virion proteins separated by SDS-PAGE identified presence of most of the predicted structural proteins. Unexpectedly, three high molecular weight subunits of phage RNA polymerase^9^ (gene products 47, 49 and 50) were also detected as part of virion structure (Fig. 2a).

**Figure 1.**
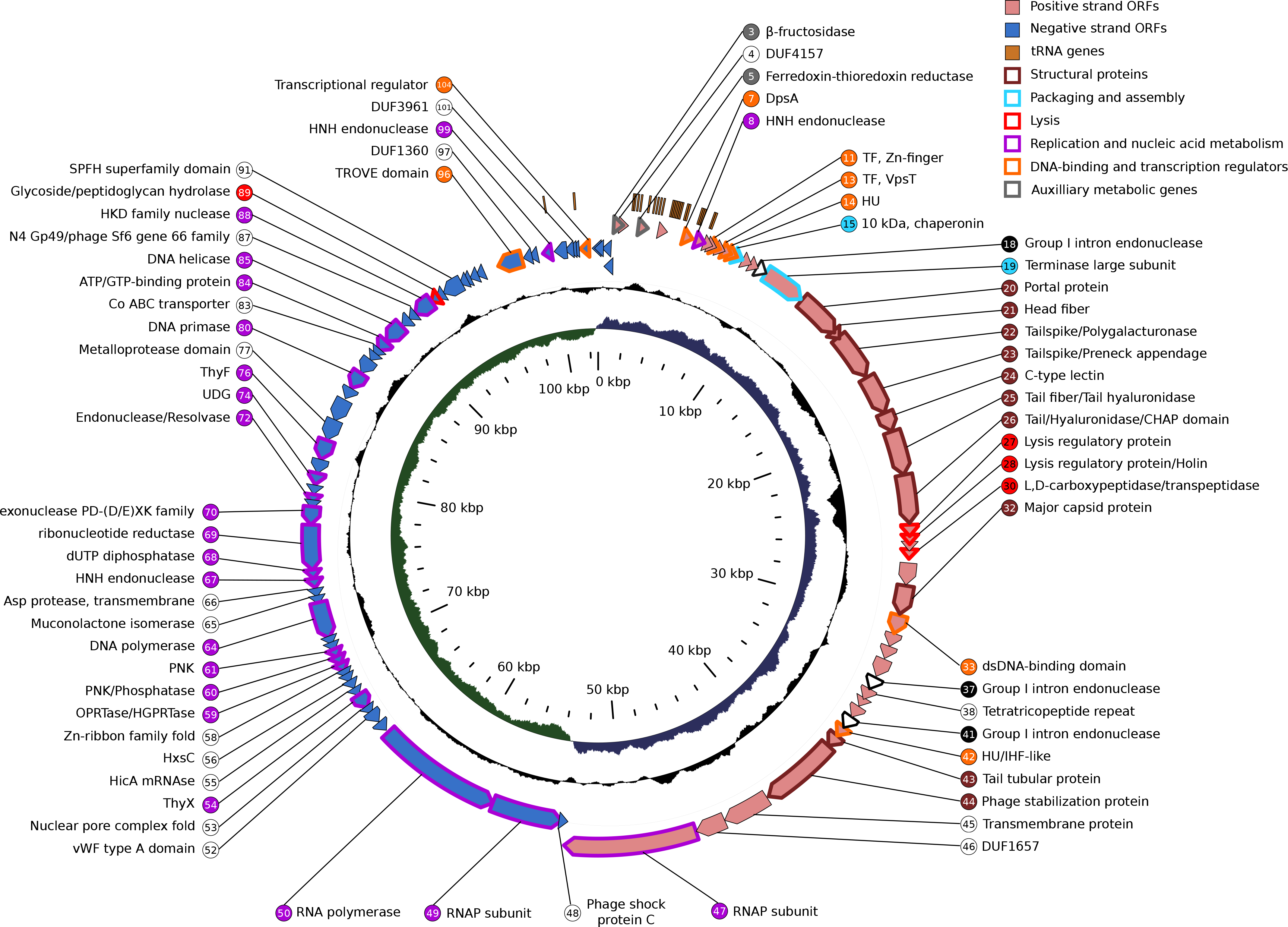
Circular map of φcrAss001 genome. Inner circle (green and blue), GC skew; Middle circle (black), G+C content; Outer circle (Pink and purple fill), protein-coding genes (ORFs); Outermost circle (orange fill), tRNA genes. Stroke colour on ORFs and fill colour on gene numbers corresponds with the general predicted function (see colour legend for details); black stroke, introns; genes with no functional annotations (coding for hypothetical proteins) are left unlabelled.

**Figure 2.**
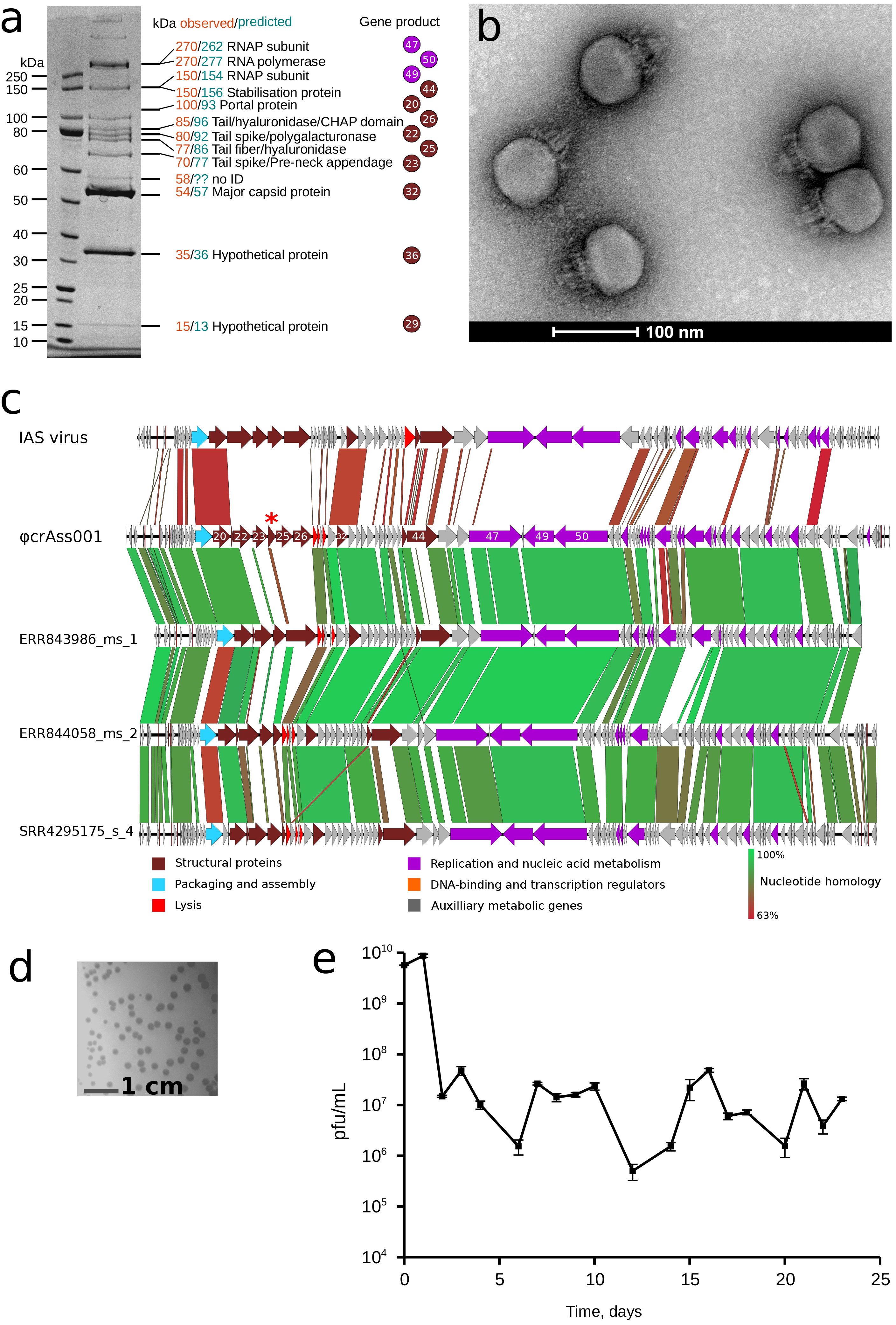
Morphology, growth and adsorption of φcrAss001. **A**, SDS-PAGE analysis of protein content in φcrAss001 virions and identification of selected polypeptides using MALDI-TOF (see methods for details); **B**, TEM image of uranyl acetate negatively contrasted φcrAss001 virions (62,000x magnification, accelerating voltage of 120kV); **C,** BLASTn comparisons between the genome of the uncultured human faecal IAS virus, φcrAss001, and uncultured human phage genomic contigs ERR843986_ms_1, ERR844058_ms_2, SRR4295175_s_4 (for contigs description see Guerin *et al.,* 2018^1^), highly variable region is marked with a red asterisk; **D**, plaque morphology of φcrAss001 after 48 h incubation in a 0.3% FAA agar overlay with *B. intestinalis* 919/174 as host strain; **E**, persistence of φcrAss001 in a periodic culture of phage infected *B*.intestinalis 919/174 with daily (or bi-daily) transfers for 23 days (see supplementary methods for details).

TEM (transmission electron microscopy) of φcrAss001 virions revealed a podoviral morphology (Fig. 2b). Phage heads are isometric with a diameter of 77.2±3.3 nm (mean±SD). Tails are 36.1±3.6 nm long with elaborate structural features and several side appendages of variable length. This is in agreement with our previous TEM observations of podoviruses of similar dimensions (~76.5 nm in diameter) from a faecal sample rich in a mixture of several highly prevalent crAss-like bacteriophages^11^.

According to the recently proposed classification scheme based on functional gene repertoire and protein sequence homology^9^, φcrAss001 fits into the IAS virus subgroup^15^ of crAss-like bacteriophages (Fig. 2c). Our recent analysis identified a common presence of similar uncultured bacteriophages in the gut microbiotas of healthy Irish and US adults, Irish elderly people, as well as in healthy Irish infants and healthy and malnourished Malawian infants^11^, where they were detectable in 62%, 44%, 50%, 10%, 14% and 16% of cases, respectively, and in some case represented up to 61% of virome reads (Fig. S1).

When genomes of φcrAss001, IAS virus and other related putative phages were compared, most of the sequence variability (including variation in the number and size of ORFs) was concentrated in a region putatively coding for tail spike/tail fiber subunits (gene products 22, 23, 25, 26, Fig. 2c). This suggests high level of variability of receptor-binding proteins and potentially high level of host specialization of crAss-like phages.

ΦcrAss001 could be effectively propagated *in vitro* using standard techniques using an EPS-producing *B. intestinalis* 919/174 as its host. It was able to form readily visible plaques in agar overlays (Fig. 2d) and reached 10^10^ pfu/mL when propagated in broth culture. In a one-step growth experiment with multiplicity of infection (MOI) of ~1 the phage demonstrated a long latent period of 120 min that was followed with a very small burst of progeny (2.5 pfu per infected cell). A second burst of roughly the same size occurred 90 min later (Fig. S2a). An adsorption curve shows that ~74% of phage bound to cells in first 5 min, and >90% were bound by 20 min (Fig. S2b). Efficiency of lysogeny/mutation rate tests demonstrated 2±1% of cells of the strain 919/174 are resistant to φcrAss001 on initial contact. These clones (potentially lysogens) were initially PCR positive for presence of φcrAss001 (using gene 20 as a target) and were resistant to the phage in plaque/spot assays. Successive rounds of propagation resulted in loss of the φcrAss001 PCR positive outcome, but the phage-resistance phenotype was retained in all clones.

Infection of exponentially growing cells of *B. intestinalis* 919/174 with φcrAss001 at different multiplicity of infections (MOI) did not result in complete culture lysis, but caused a delay in stationary phase onset time and final density at stationary phase (Fig. S2c). A brief lysis period occurred in the first few hours after infection, with timing dependent on the MOI, followed by recovery of bacterial growth. In order to investigate the fate of phage in co-culture, bacterial cells infected with phage at high MOI were allowed to reach stationary phase and then passaged daily or bi-daily for a period of 23 days (Fig. 2e). Phage titre, which on the first two passages reached ~10^10^, was reduced and remained steady between 10^6^−10^8^ for the duration of the experiment.

Collectively, this suggests that φcrAss001 uses an unusual infection strategy to replicate *in vitro* on its *B. intestinalis* host very efficiently on semi-solid agar (giving rise to large clear plaques) and yet propagate in liquid culture without causing lysis of the host bacterium. Despite not being able to form true lysogens or pseudolysogens, the phage seem to be able to co-exist with its host in an equilibrium which is likely to confer an ecological advantage to both partners (similar to the recently described carrier state life cycle^19^). Taken together, these results are puzzling. Given that in liquid culture we used an MOI of 1 and there is a high adsorption rate, we cannot explain why two bursts seemed to occur and why the broth culture did not clear. We can conclude that the phage probably causes a successful lytic infection in a subset of infected cells (giving rise to a ‘false’ overall burst size of 2.5), but must enter into an alternative interaction with some or all of the remaining cells. Overall, this allows both phage and host to co-exist in a stable interaction over prolonged passages. This may explain why the *Bacteroides* host and crAssphage can both remain in high numbers for pronged period in the gut. The nature of this interaction warrants further investigation.

Published studies of gut viromes in humans and germ-free mice with transplanted human viromes^12,20^, as well as our own unpublished longitudinal observations of the human gut virome support the hypothesis that crAssphage use an unusual strategy to establish themselves at high levels in the gut and to then persist stably within the microbial communities for several weeks to as long as several months or even years. The absence of any detectable integrase gene and our inability to isolate stable lysogens, together with the lack of any evidence that this or any other crAss-like bacteriophages can occur in the form of prophages, suggest that pseudolysogeny and other mechanisms, such as physical binding of phage to the intestinal mucous gel^21^ might be responsible for long term persistence of crAss-like phages *in vivo.* Further studies will be required to fully understand the replication cycle of crAss-like bacteriophages, their peculiar ability to persist at high titres in the human gut microbiota, as well as their significance for human intestinal physiology and disease. The first report of a phage-host pair should accelerate our understanding of these highly abundant and unusual phages.

## End notes

## Acknowledgements

This research was conducted with the financial support of Science Foundation Ireland (SFI) under Grant Number SFI/12/RC/2273 a Science Foundation Ireland’s Spokes Programme which is co-funded under the European Regional Development Fund under Grant Number SFI/14/SP APC/B3032, and a research grant from Janssen Biotech, Inc.

**Supplementary Information** is available in the online version of the paper (Supplementary table 1, Supplementary figure 1, Supplementary figure 2, Supplementary methods)

## Author contributions

ANS, EVK and CBF performed the experiments; ANS, SRS, LAD, RPR and CH analysed the data; RPR and CH supervised the project.

## Author information

Reprints and permissions information is available at www.nature.com/reprints. The authors declare no competeng financial interests. Correspondence and requests for materials should be addressed to CH or ANS.

